# Cell clusters adopt a collective amoeboid mode of migration in confined non-adhesive environments

**DOI:** 10.1101/2020.05.28.106203

**Authors:** Diane-Laure Pagès, Emmanuel Dornier, Jean De Seze, Li Wang, Rui Luan, Jérôme Cartry, Charlotte Canet-Jourdan, Joel Raingeaud, Raphael Voituriez, Mathieu Coppey, Matthieu Piel, Fanny Jaulin

## Abstract

Cell migration is essential to most living organisms. Single cell migration involves two distinct mechanisms, either a focal adhesion- and traction-dependent mesenchymal motility or an adhesion-independent but contractility-driven propulsive amoeboid locomotion. Cohesive migration of a group of cells, also called collective cell migration, has been only described as an adhesion- and traction-dependent mode of locomotion where the driving forces are mostly exerted at the front by leader cells. Here, by studying primary cancer specimens and cell lines from colorectal cancer, we demonstrate the existence of a second mode of collective migration which does not require adhesion to the surroundings and relies on a polarised supracellular contractility. Cell clusters confined into non-adhesive microchannels migrate in a rounded morphology, independently of the formation of focal adhesions or protruding leader cells, and lacking internal flow of cells, ruling-out classical traction-driven collective migration. Like single cells migrating in an amoeboid fashion, the clusters display a supracellular actin cortex with myosin II enriched at the rear. Using pharmacological inhibitors and optogenetics, we show that this polarised actomyosin activity powers migration and propels the clusters. This new mode of migration, that we named collective amoeboid, could be enabled by intrinsic or extrinsic neoplasic features to enable the metastatic spread of cancers.

**One Sentence Summary:** Clusters organise as polarised and contractile super-cells to migrate without adhesion.

## Main Text

Migration is a fundamental property of cells. Emerging in early eukaryotes, migration supports individual cell displacement as well as metazoan development and homeostasis (*1*). It is also deregulated in pathological conditions, such as cancer, where it fuels their metastatic spread (*2*). Two distinct mechanisms are used by single cells to generate the migration forces (*3*). They result from the cells’ ability to adhere, or not, to the surrounding extracellular matrix (ECM) and their level of contractility. In traction-based mesenchymal migration, integrin interaction with the ECM and focal adhesion formation convert branched-actin polymerisation into large protrusions and forward forces (*3, 4*). In contrast, amoeboid single cells use a propulsive locomotion that does not require specific adhesion and is driven by acto-myosin contractility of the rear (*5, 6*). Cells can also move in a cohesive manner as a group (7–9). At the front of the cluster, leader cells form prominent protrusions where the combined action of actin polymerization and integrin engagement triggers lamellipodia and focal adhesion formation. Using the substrate as an anchor, leaders pull on follower cells, instructing directionality and generating important traction forces (*10, 11*). The contribution of follower cells is more elusive, but it has recently been shown that their increased contractility produces a treadmilling of lateral cells to support the migration of neural crest clusters (*12*). To date, our knowledge on collective cell movement suggests it only takes the form of an adhesion-dependent traction-based mode of locomotion and whether it could also occur through an alternative mechanism has not been investigated.

Through the analysis of primary tumour explants retrieved from patients with metastatic colorectal cancer (CRC), we identified TSIPs (Tumour Spheres with Inverted Polarity) as tumour cell clusters with an inverted apico-basolateral polarity (*13*). The atypical topology of the clusters exposes carcinoma cells’ apical membranes to the microenvironment and precludes adhesion receptors, such as integrins, to interact with the surrounding ECM in the peritumoral stroma (Fig. 1A). Yet, TSIPs efficiently invade tissues and are associated with high metastatic burden and poor patient prognosis (*13*). This suggests that the motility of TSIPs does not require integrin function and raises the possibility of an adhesion-independent mode of collective cell migration. To test this hypothesis, we engineered micro-devices (channels and chambers) deprived of any physiological substrates and chemotactic cues by coating them with the anti-adhesive polymer polyethylene glycol (PEG, Fig. 1B). Time-lapse imaging proved that TSIPs obtained from two independent patients migrated into the non-adhesive microchannels (Fig. 1C(a), fig. S1A and movie S1). To determine whether this type of migration is specific to TSIPs or could be used by other collectives, we assessed the migration of clusters assembled from colorectal carcinoma cell lines in PEG-coated microchannels. Indeed, clusters from HT29, HT29-MTX and three lines of circulating tumour cells (CTC31,44, 45, (*14*)) were able to collectively migrate under these conditions (Fig. 1, C(b) to E, and movie S2). To characterise this new mode of collective migration, considering the durations of consecutive migration and pauses (fig. S1, B and C), we monitored trajectories of individual clusters every hour over one day (Fig. 1D). Some clusters did not move in the course of the experiment while some reached-up to 2 mm/d. Average speeds ranged from 150±21μm/d to 77±4μm/d for TSIPs and HT29s which are the fastest. The migration of CTCs was slower, varying from 37±2μm/d to 27±2μm/d (Fig. 1E) (values are expressed as speed ± standard error of the mean). Clusters displayed a very persistent migration over time (from 0.65±0.03 to 0.75±0.03 in average) and could reach a maximum instantaneous speed of 28±3μm/h in average (Fig. 1D and F, and fig. S1D). Although quite slow when compared with single cell migration in experimental settings, this is in the order of magnitude of collective migration speeds reported in vivo (*15, 16*).

**Fig. 1.**
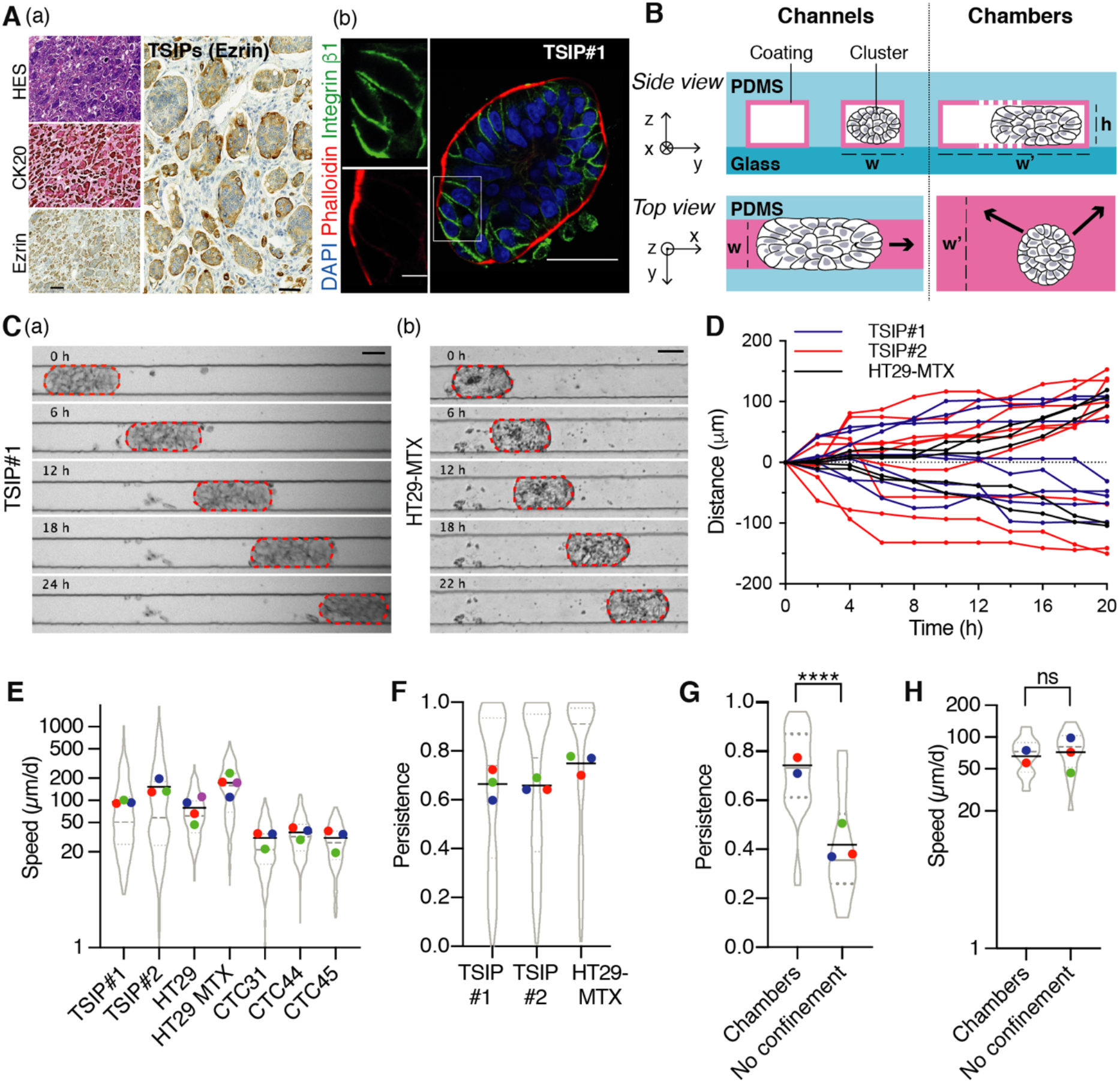
Cell clusters migrate into non-adhesive microchannels. (**A**) Representative tumour specimen of CRC (micropapillary subtype) stained with Hematoxylin/Eosin/Saffron (HES), anti-Cytokeratin 20 (CK20) and anti-Ezrin at low (left) and high (right) magnification (a) and immunofluorescence of a **t**ransverse section of TSIP#I embedded in collagen-1 for 24h (b). The boxed region is shown at high magnification. (**B**) Schematic representation, not to scale, of the channels (width: w=60*μ*m; height: h=30*μ*m) (left) and chambers (width: w’=500*μ*m) (right). (**C**) Time-lapse sequences of migration of TSIP#I (a) and HT29-MTX (b) in PEG-coated microchannels. (**D**) Representative tracks of clusters migrating in one direction (positive numbers) or the other (negative numbers) in PEG-coated microchannels. n=5 to 9 representative clusters of each cell type. (E and F) Clusters’ migration speed (E, log2-scale, n=118 to 208 clusters per cell type) and persistence (F, n=107 to 124 clusters per cell type) in PEG-coated microchannels over one day. (**G** and **H**) Speed (G, log2-scale, unpaired Student’s t-test) and persistence (H, Mann-Whitney test) for HT29-MTX migration over a day, depending on their confinement. Designs are coated with PEG+F127. n=30 clusters (chambers), n=18 clusters (no confinement), ns, not significant, ****P<0.0001 (Kruskal-Wallis test). Violin plots display the whole population of clusters, median (dashed grey line) and quartiles (dotted grey lines). The coloured dots represent the mean of each independent experiments and the black line the mean of all experiments for each condition. All data represented as violin plots are from N=3 independent experiments, except HT29 and HT29-MTX in (E, N=4) and chambers in (G) and (H) (N=2). Scale bars: 200*μ*m (A,a, low magnification), 50*μ*m [(A,a, high magnification), (A,b, low magnification), (C)] and 10*μ*m (A,b, high magnification).

We next assessed the role of confinement by comparing migration of clusters confined in one dimension (microchambers, Fig. 1B and fig. S2A) or not confined (loading chambers, fig. S2B). Confinement did not increase clusters’ speed but favoured persistence, as described for single cells (Fig. 1, G and H) (*6, 17, 18*). Once confined into microchannels, small clusters migrate as fast as the largest ones, showing no correlation between speed and size (fig. S2, C to E). In all instances, the collective migration is associated with a compact rounded morphology into non-adhesive microchannels that contrasts with the loose and spread shape clusters can adopt when the microchannels are coated with collagen-1 (Fig. 2A). Measuring the contact angles between the cluster boundary and the microchannel walls highlights the “dewetting” morphology of the clusters and the absence of protrusion for HT29, HT29-MTX and TSIPs migrating in PEG-coated microchannels (Fig. 2, A and B). Collagen-1 coating reduces HT29 and HT29-MTX migration speed while, as expected, TSIPs remain unaffected due to their inverted polarity (Fig. 2C and 1A). Together, these experiments suggest that confined cell clusters can display a persistent motility in non-adhesive environments.

**Fig. 2.**
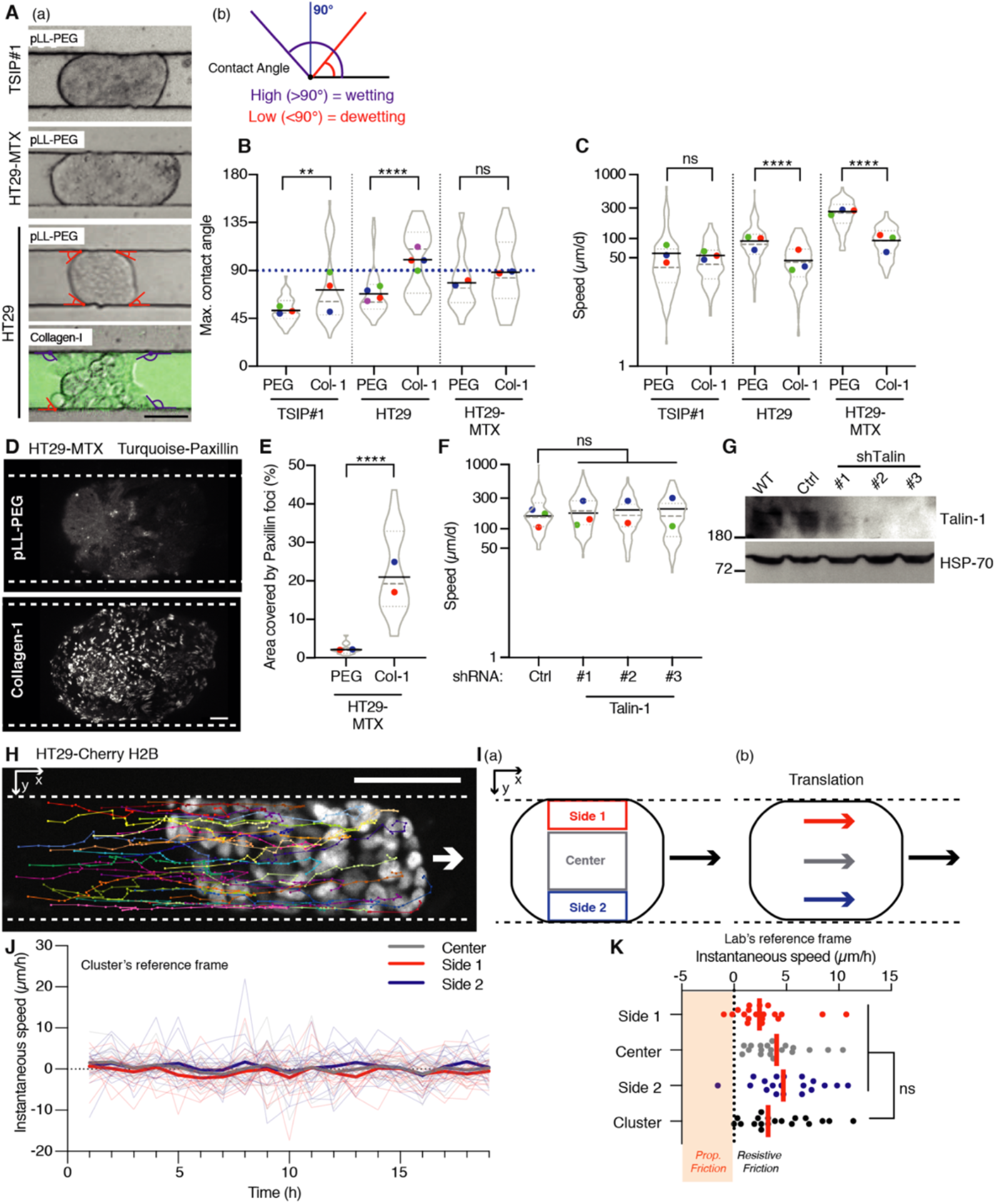
Collective migration occurs without focal-adhesions nor cell flows. (**A** to **C**) Representative images (A), maximum contact angles (B) and speeds (C) of clusters migrating in PEG- or Cy5-collagen-1(Col-1)-coated microchannels. Wetting or dewetting contact angles are represented on the cluster (A,a) and as a schematic representation (A,b); n=39 to 59 clusters were used to quantify the maximum contact angle (B, Mann Whitney test). n=9O to 166 clusters were analysed to calculate migration speeds (C, two-tailed Student’s t test). (**D** and **E**) Representative images of the bottom plan of the cluster (D) and area covered by paxillin foci (E) for HT29-MTX stably expressing mTurquoise-Paxillin in PEG- or Col-1-coated microchannels. n=19 to 28 clusters (E, Mann-Whitney test). (**F** and **G**) Speed of clusters in PEG-coated microchannels (F) and corresponding western blot (G) for HT29-MTX stably expressing 3 different shRNA targeting Talin-1 or a control shRNA. n=62 to 154 clusters (F, one-way ANOVA). (**H**) Representative nuclei tracks of the middle cross-section of an HT-29 cluster stably expressing mCherry-H2B, migrating one day in a PEG-coated microchannel. (**I**) Definition of areas in the cluster (a), to scale, and schematic representation of cluster’s global translation (b). Side 1: cluster’s side going most rearward/less forward in average. (**J**) Instantaneous speed of cells in each area, in the reference frame of the clusters. Bold lines: average of individuals tracks calculated from regions of n=16 clusters from 3 independent experiments. (**K**) For one representative cluster, instantaneous speed of cells in each area and cluster speed, lab’s reference frame. Orange: cells could generate propulsive (Prop.) friction. n=19 timepoints (paired Friedman test). Experiments were performed independently for each cell lines. Violin plots are described in Fig. 1 legend and coloured dots refer to the same experiment. All data represented as violin plots are from N=3 independent experiments, except HT29-MTX in (B) and (E), and sRNA#2 and #3 in (F) that were performed twice, ns, not significant, **P<0.01, ****P<0.0001. Scale bars: 50μm [(A) and (H)J, 10*μ*m (D).

To directly study the contribution of focal adhesion to cluster migration, we expressed turquoise-tagged paxillin in HT29-MTX. While fluorescent paxillin revealed numerous foci at collagen-1 interface, they were nearly absent in PEG-coated microchannels (Fig. 2, D and E, and movie S3). To functionally assess the participation of focal adhesions, we silenced talin-1, an essential component of integrin-mediated functions (*19*). This had no effect on HT29-MTX cluster migration (Fig. 2, F and G). Hence, interfering with intrinsic and extrinsic components of cell adhesion to their substrate demonstrates that clusters can migrate without the conventional molecular machinery powering traction-based collective migration (*20–22*).

We next tested whether a coordinated retrograde flow of cells, or cell treadmilling, could participate in this non-adhesive cluster migration, as was proposed before in a developmental context (*12*). To this end, we expressed cherry-tagged histone 2B (H2B) in HT29 clusters to monitor individual cell movements during migration in PEG-coated channels. Confocal fluorescence imaging showed that individual cell tracks follow the trajectory of the centre of mass of the cluster (Fig. 2H, fig. S3A and movie S4). To precisely test the hypothesis of flows of cells generating migration (Fig. S3, B to G), three areas were defined on a middle cross section of the cluster and cell trajectories were examined (Fig. 2I(a)). In each area, the measurement of the instantaneous speed of cells in the cluster’s reference frame directly confirmed the absence of significant fluxes of cells (Fig. 2J and fig. S3, C and F). As a consequence, the relative velocity of cells in contact with the microchannel walls was positive, in the direction of the motion of the cluster (Fig. 2K). In the hypothesis of a propulsion mechanism based on flows of cells, this would indicate the presence of resistive friction forces only and the absence of propulsion forces, excluding a mechanism based on an internal treadmilling of cells (fig. S3 D and G). Thus, clusters translate, with all cells remaining at the same relative position in the group during migration (Fig. 2I(b) and fig.S3A).

The classical mechanisms of traction-driven collective migration being ruled-out, we reasoned that the acto-myosin cytoskeleton could power focal adhesion-independent cluster migration, as it does in amoeboid single cells (*5, 6, 23, 24*). Expressing the fluorescent probes F-tractin and myosin light chain (MLC) in HT29-MTX revealed a robust peripheral supracellular acto-myosin cortex as reported at the boundaries of other migrating collectives (Fig. 3A) (*25*). This peripheral cortex is evenly distributed in static clusters, however, during migration, it exhibits a front/rear polarisation, with a 1.5- and 1.8-fold enrichment toward the back of the cluster for F-tractin and MLC respectively (Fig. 3, A and B, and movie S5). To assess whether this supracellular acto-myosin cortex contributes to cluster migration, we first used pharmacological inhibitors. Interfering with Myosin-II or ROCK activities using blebbistatin and Y27632 reduced TSIP#1 migration speed from 80±7μm/d to 50±6 μm/d and 43±3 μm/d, respectively (Fig. 3, C and D, and fig. S4A). Similarly, using these inhibitors on HT29-MTX decreased their migration by 3- and 2.9-fold, respectively (Fig. 3, C and D, and fig. S4B). Then, we tested whether increasing contractility at the rear was sufficient to power cluster migration. To this end, we used optogenetics to manipulate acto-myosin contractility via its upstream regulator RhoA. We infected HT29-MTX cells with the optoRhoA system, which enables an acute spatiotemporal recruitment of RhoA activator ARHGEF11 to the membrane using the CRY2/CIBN light gated optogenetic dimerization system (fig. S5, A and B) (*26*). We illuminated one side of the clusters, either at the front of already moving clusters or randomly for static ones. We monitored their trajectories for up to 20 hours by using an automated stage and activation routine maintaining a constant illumination region despite the movement of the cluster (Fig. 3E). Control clusters expressing the optogenetic dimeriser without RhoA activator pursued their migration in their initial direction with their speed sometimes reduced by mild phototoxicity or large local protein recruitment at the membrane (*27*). In contrast, illuminating clusters expressing optoRhoA impacted their migratory behaviours. Light stimulation initiated the migration of static clusters and was even able to revert the direction of migrating ones (Fig. 3, F to H, and fig. S5C, and movie S6, 7). This demonstrates that increasing acto-myosin activity in a subset of cells is both sufficient to induce migration and dictate directionality taken by the entire cluster (Fig. 3, F to H, and fig. S5C). Altogether, these experiments show that a supracellular polarised acto-myosin contractility contributes to the driving force that propels clusters of cells into non-adhesive environments.

**Fig. 3.**
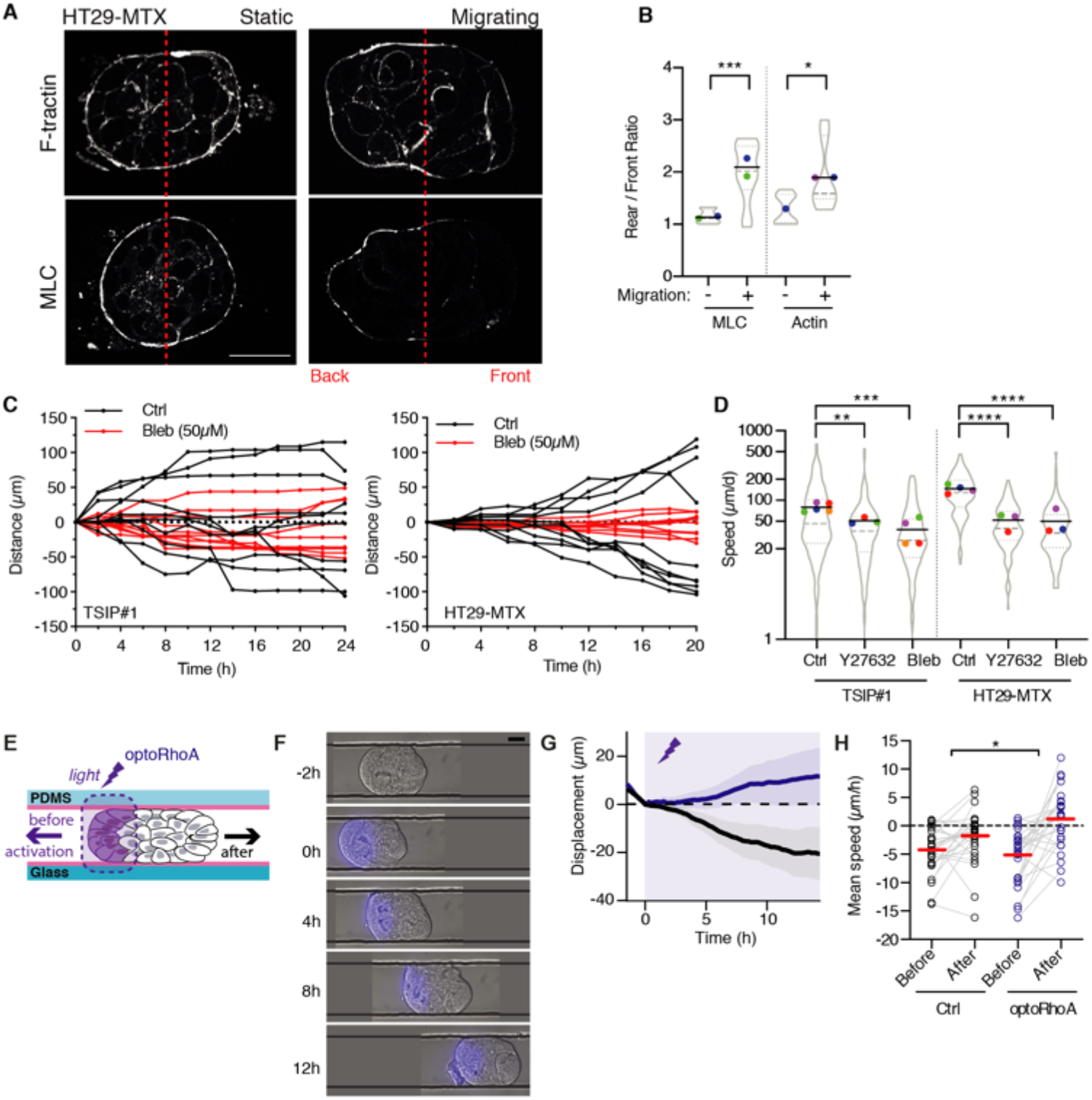
A supracellular contractile actin cortex powers focal adhesion-independent collective migration. (**A**) Median section of HT29-MTX stably expressing F-tractin-mRuby3 (top) and mTurquoise-MLC (bottom), in PEG+F127-coated channels. (**B**) Quantification of the F-tractin (Actin) and MLC (Myosin Light Chain) ratios between rear and front of the clusters, as indicated in Methods, n=7 to 13 clusters (Mann-Whitney test). (**C**) Representative tracks of clusters treated with Blebbistatin. n=10 clusters each. (**D**) Speed of clusters treated with Y27632, Blebbistatin, or DMSO (Ctrl), log2-scale. n=106 to 205 clusters (Mann-Whitney test). (**E** to **H**) Optogenetic manipulations: Experimental setup (E), representative time-lapse sequences (F, dark grey lines are microchannel walls), migration profiles (G, means ± sem) and mean speeds before (−2h<t<0h) and after (4h<t<10h) optogenetic activation (H). In G and H, the control (CRY2PHR-mCherryN1/CIBN-eGFP-CaaX) is represented as black line or dots and optoRhoA as purple line or dots. The purple zones represent the optogenetic activation period, n=27 clusters from 3 independent experiments (two-tailed Student’s t test on the after-before speed differences between control and optogenetic constructs). Experiments were performed independently for each cell types. Violin plots are described in Fig. 1 legend and coloured dots refer to the same experiment. All data represented as violin plots are from N=3 independent experiments, except in (B, N=2, and N=1 for Actin in non-migrating clusters), and TSIP#I in Blebbistatin (E) (N=4). ns, not significant, *P<0.05, “P<0.01, ***P<0.001, **** P<0.0001. Scale bars, 30 *μ*m.

Here, we report a second mode of collective migration that shares striking features with amoeboid single cell motility (Fig. 4). Cell clusters collectively migrate within non-adhesive microchannels in the absence of protruding leader cells, focal adhesions or cell flows. Similar to amoeboid cells, clusters mobilise the contractility of a supracellular acto-myosin cortex at the rear and adopt a propulsive mode of migration. By analogy, we named this new mode of cell locomotion “collective amoeboid migration”. Tumour cells have the capacity to hijack the 3 modes of cell migration described to date (*28*). Here, we show that colorectal cancer primary specimens and cell lines can also adopt collective amoeboid migration. This results from intrinsic oncogenic features, such as in TSIPs, where the inverted apico-basolateral polarity prevents cluster adhesion to ECM-rich tissues (*13*). This propulsion-based mode of collective migration could also be enabled as a non-cell autonomous process, when cancer cell clusters are exposed to environments deprived of conventional ECM. These encompass major dissemination routes such as the lumen of lymphatic vessels or the peritoneal and pleural cavities (*29–33*). Collective amoeboid migration could thus foster cancer metastatic spread.

**Fig. 4.**
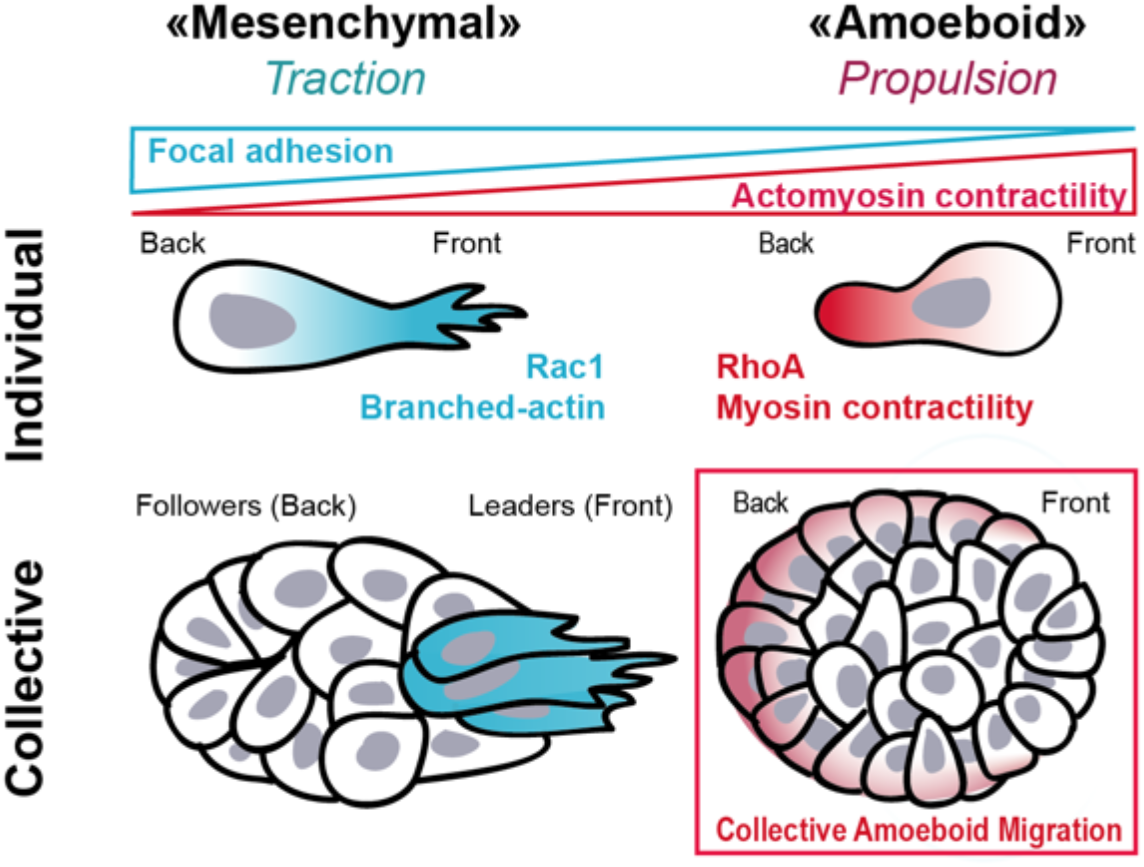
Model of cell migration modes. Schematic representation of the different modes of cell migration based on the ability of the cells to adhere to each other (individual versus collective migration) and to their environment (mesenchymal versus amoeboid). This results in traction- or propulsion-based locomotion, where the force is respectively generated by Rad-dependent actin polymerization or RhoA-dependent actomyosin contractility. The second mode of collective migration we report here occurs without the formation of focal adhesion and relies on acto-myosin contractility at the rear. By analogy with single cells, it has been named «collective amoeboid migration» and represents a fourth mode of cell migration.

## Acknowledgements

We thank the members of the Jaulin, Piel and Montagnac laboratories for helpful discussions. We thank T. Manoliu and C. Laplace from the PFIC facility for their technical help. We thank M. Polrot from the PFEP facility for assistance with the animal work. We thank the C. Robert lab for the plasma cleaner. We thank J. Pannequin for the CTC cell lines. We thank W. Beng and K. Vaidziulyte for the automated tracking algorithm used in optogenetics. Funding: This work was supported by grants from INCA-PLBIO (2018-1-PLBIO-04-IGR-1), ARC (PAJ20181208275) and LNCC-IDF (R19102LL) to F.J. as well as fund raising against colorectal cancer from the Gustave Roussy foundation. M.C. thanks the Labex CelTisPhyBio (ANR-10-LBX-0038) and the Institut Convergences Q-life ANR-17-CONV-0005. We thank the Philantropia fellowship to D.-L.P., the FRM fellowship to E.D., the AMX fellowship to J.D.S. and the Taxe d’Apprentissage 2019 (to D.-L.P., Univ. Paris-Saclay, France).

## Author contributions

All authors provided intellectual input. D.-L.P., E.D., and R.L. designed the research, performed the experiments and analysed the data. J.D.S. and M.C. conceived, performed and analysed the optogenetics experiments. L.W. and M.P. designed and validated the microfabricated devices. J.C. prepared the organoids and helped for the animal work. C.C.-J. performed immunostainings on TSIPs. J.R. analysed the contractility of cell clusters. C.M.C, R.V. and M.P. provided scientific input. F.J. conceptualised the project, designed the research and wrote the manuscript.

## Data and materials availability

All data is available in the main text or the supplementary materials.

## References and Notes

1. K. M. Yamada, M. Sixt, Mechanisms of 3D cell migration. Nat Rev Mol Cell Biol. 20, 738–752 (2019).

2. A. W. Lambert, D. R. Pattabiraman, R. A. Weinberg, Emerging Biological Principles of Metastasis. Cell. 168, 670–691 (2017).

3. D. L. Bodor, W. Pönisch, R. G. Endres, E. K. Paluch, Of Cell Shapes and Motion: The Physical Basis of Animal Cell Migration. Dev. Cell. 52, 550–562 (2020).

4. M. Innocenti, New insights into the formation and the function of lamellipodia and ruffles in mesenchymal cell migration. Cell Adh Migr. 12, 401–416 (2018).

5. E. K. Paluch, I. M. Aspalter, M. Sixt, Focal Adhesion-Independent Cell Migration. Annu. Rev. Cell Dev. Biol. 32, 469–490 (2016).

6. Y.-J. Liu, M. Le Berre, F. Lautenschlaeger, P. Maiuri, A. Callan-Jones, M. Heuzé, T. Takaki, R. Voituriez, M. Piel, Confinement and low adhesion induce fast amoeboid migration of slow mesenchymal cells. Cell. 160, 659–672 (2015).

7. P. Friedl, J. Locker, E. Sahai, J. E. Segall, Classifying collective cancer cell invasion. Nat. Cell Biol. 14, 777–783 (2012).

8. P. Friedl, D. Gilmour, Collective cell migration in morphogenesis, regeneration and cancer. Nat. Rev. Mol. Cell Biol. 10, 445–457 (2009).

9. K. J. Cheung, A. J. Ewald, A collective route to metastasis: Seeding by tumor cell clusters. Science. 352, 167–169 (2016).

10. R. Mayor, S. Etienne-Manneville, The front and rear of collective cell migration. Nat. Rev. Mol. Cell Biol. 17, 97–109 (2016).

11. E. Theveneau, C. Linker, Leaders in collective migration: are front cells really endowed with a particular set of skills? F1OOORes. 6, 1899 (2017).

12. A. Shellard, A. Szabó, X. Trepat, R. Mayor, Supracellular contraction at the rear of neural crest cell groups drives collective chemotaxis. Science. 362, 339–343 (2018).

13. O. Zajac, J. Raingeaud, F. Libanje, C. Lefebvre, D. Sabino, I. Martins, P. Roy, C. Benatar, C. Canet-Jourdan, P. Azorin, M. Polrot, P. Gonin, S. Benbarche, S. Souquere, G. Pierron, D. Nowak, L. Bigot, M. Ducreux, D. Malka, C. Lobry, J.-Y. Scoazec, C. Eveno, M. Pocard, J.-L. Perfettini, D. Elias, P. Dartigues, D. Goéré, F. Jaulin, Tumour spheres with inverted polarity drive the formation of peritoneal metastases in patients with hypermethylated colorectal carcinomas. Nature Cell Biology, 1 (2018).

14. F. Grillet, E. Bayet, O. Villeronce, L. Zappia, E. L. Lagerqvist, S. Lunke, E. Charafe-Jauffret, K. Pham, C. Molck, N. Rolland, J. F. Bourgaux, M. Prudhomme, C. Philippe, S. Bravo, J. C. Boyer, L. Canterel-Thouennon, G. R. Taylor, A. Hsu, J. M. Pascussi, F. Hollande, J. Pannequin, Circulating tumour cells from patients with colorectal cancer have cancer stem cell hallmarks in ex vivo culture. Gut. 66, 1802–1810 (2017).

15. D. Krndija, F. El Marjou, B. Guirao, S. Richon, O. Leroy, Y. Bellaiche, E. Hannezo, D. Matic Vignjevic, Active cell migration is critical for steady-state epithelial turnover in the gut. Science. 365, 705–710 (2019).

16. B. Weigelin, G.-J. Bakker, P. Friedl, Intravital third harmonic generation microscopy of collective melanoma cell invasion: Principles of interface guidance and microvesicle dynamics. Intravital. 1, 32–43 (2012).

17. B. Winkler, I. S. Aranson, F. Ziebert, Confinement and substrate topography control cell migration in a 3D computational model. Commun Phys. 2, 1–11 (2019).

18. J. A. Mosier, A. Rahman-Zaman, M. R. Zanotelli, J. A. VanderBurgh, F. Bordeleau, B. D. Hoffman, C. A. Reinhart-King, Extent of Cell Confinement in Microtracks Affects Speed and Results in Differential Matrix Strains. Biophys. J. 117, 1692–1701 (2019).

19. Z. Sun, M. Costell, R. Fässler, Integrin activation by talin, kindlin and mechanical forces. Nat. Cell Biol. 21, 25–31 (2019).

20. F. Gunawan, A. Gentile, R. Fukuda, A. T. Tsedeke, V. Jiménez-Amilburu, R. Ramadass, A. Iida, A. Sehara-Fujisawa, D. Y. R. Stainier, Focal adhesions are essential to drive zebrafish heart valve morphogenesis. J Cell Biol. 218, 1039–1054 (2019).

21. E. H. Barriga, K. Franze, G. Charras, R. Mayor, Tissue stiffening coordinates morphogenesis by triggering collective cell migration in vivo. Nature. 554, 523–527 (2018).

22. Y. Hegerfeldt, M. Tusch, E.-B. Bröcker, P. Friedl, Collective cell movement in primary melanoma explants: plasticity of cell-cell interaction, beta1-integrin function, and migration strategies. Cancer Res. 62, 2125–2130 (2002).

23. T. Lämmermann, B. L. Bader, S. J. Monkley, T. Worbs, R. Wedlich-Söldner, K. Hirsch, M. Keller, R. Förster, D. R. Critchley, R. Fässler, M. Sixt, Rapid leukocyte migration by integrin-independent flowing and squeezing. Nature. 453, 51–55 (2008).

24. M. Bergert, A. Erzberger, R. A. Desai, I. M. Aspalter, A. C. Oates, G. Charras, G. Salbreux, E. K. Paluch, Force transmission during adhesion-independent migration. Nat. Cell Biol. 17, 524–529 (2015).

25. A. Shellard, R. Mayor, Supracellular migration - beyond collective cell migration. J. Cell. Sci. 132 (2019), doi:10.1242/jcs.226142.

26. L. Valon, A. Marín-Llauradó, T. Wyatt, G. Charras, X. Trepat, Optogenetic control of cellular forces and mechanotransduction. Nat Commun. 8, 14396 (2017).

27. X. Meshik, P. R. O’Neill, N. Gautam, Physical Plasma Membrane Perturbation Using Subcellular Optogenetics Drives Integrin-Activated Cell Migration. ACS Synth Biol. 8, 498–510 (2019).

28. P. Friedl, S. Alexander, Cancer invasion and the microenvironment: plasticity and reciprocity. Cell. 147, 992–1009 (2011).

29. D. J. Ruiter, J. H. van Krieken, G. N. van Muijen, R. M. de Waal, Tumour metastasis: is tissue an issue? Lancet Oncol. 2, 109–112 (2001).

30. S.-B. Lim, C. S. Yu, S. J. Jang, T. W. Kim, J. H. Kim, J. C. Kim, Prognostic significance of lymphovascular invasion in sporadic colorectal cancer. Dis. Colon Rectum. 53, 377–384 (2010).

31. V. Barresi, L. Reggiani Bonetti, E. Vitarelli, C. Di Gregorio, M. Ponz de Leon, G. Barresi, Immunohistochemical assessment of lymphovascular invasion in stage I colorectal carcinoma: prognostic relevance and correlation with nodal micrometastases. Am. J. Surg. Pathol. 36, 66–72 (2012).

32. M. Brown, F. P. Assen, A. Leithner, J. Abe, H. Schachner, G. Asfour, Z. Bago-Horvath, J. V. Stein, P. Uhrin, M. Sixt, D. Kerjaschki, Lymph node blood vessels provide exit routes for metastatic tumor cell dissemination in mice. Science. 359, 1408–1411 (2018).

33. J. Law, E. Martin, “Transcoelomic Spread”, in Concise Medical Dictionary (Oxford University Press, ed. 10, 2020).

34. S. Julien, A. Merino-Trigo, L. Lacroix, M. Pocard, D. Goéré, P. Mariani, S. Landron, L. Bigot, F. Nemati, P. Dartigues, L. B. Weiswald, D. Lantuas, L. Morgand, E. Pham, P. Gonin, V. Dangles-Marie, B. Job, P. Dessen, A. Bruno, A. Pierré, H. De Thé, H. Soliman, M. Nunes, G. Lardier, L. Calvet, B. Demers, G. Prévost, P. Vrignaud, S. Roman-Roman, O. Duchamp, C. Berthet, Characterization of a large panel of patient-derived tumor xenografts representing the clinical heterogeneity of human colorectal cancer. Clin. Cancer Res. 18, 5314–5328 (2012).

35. M. J. Kennedy, R. M. Hughes, L. A. Peteya, J. W. Schwartz, M. D. Ehlers, C. L. Tucker, Rapid blue-light-mediated induction of protein interactions in living cells. Nat Methods. 7, 973–975 (2010).

36. K. E. Rothenberg, D. W. Scott, N. Christoforou, B. D. Hoffman. Vinculin force-sensitive dynamics at focal adhesions enable effective directed cell migration. Biophys J. 114, 1680–1694 (2018).

37. A. Hayer, L. Shao, M. Chung, L.-M. Joubert, H. W. Yang, F. C. Tsai, A. Bisaria, E. Betzig, T. Meyer. Engulfed cadherin fingers are polarized junctional structures between collectively migrating endothelial cells. Nat Cell Biol. 18, 1311–1323 (2016).

38. C. M. Kenific, S. J. Stehbens, J. Goldsmith, A. M. Leidal, N. Faure, J. Ye, T. Wittmann, J. Debnath. NBR1 enables autophagy-dependent focal adhesion turnover. J. Cell Biol. 212, 577–590 (2012).

39. J. Schindelin, I. Arganda-Carreras, E. Frise, V. Kaynig, M. Longair, T. Pietzsch, S. Preibisch, C. Rueden, S. Saalfeld, B. Schmid, J.-Y. Tinevez, D. J. White, V. Hartenstein, K. Eliceiri, P. Tomancak, A. Cardona. Fiji: an open-source platform for biological-image analysis. Nat. Methods 9, 676–682 (2012).

40. J.-Y. Tinevez, N. Perry, J. Schindelin, G. M. Hoopes, G. D. Reynolds, E. Laplantine, S. Y. Bednarek, S. L. Shorte, K. W. Eliceiri. TrackMate: An open and extensible platform for single-particle tracking. Methods 115, 80–90 (2017).

41. S. J. Lord, K. B. Velle, R. D. Mullins, L. K. Fritz-Laylin. SuperPlots: Communicating reproducibility and variability in cell biology. J. Cell Biol. 219, e202001064 (2020).

