## Supplementary Materials for "Cell clusters adopt a collective amoeboid mode of migration in confined non-adhesive environments"

### Materials and Methods

#### Biological material and cell culture

*Human primary specimens.* The human study protocols followed all relevant ethical regulations in accordance with the declaration of Helsinki principles. The study was approved by the ethic committee at Gustave Roussy hospital and written informed consent was obtained from all patients.

*Tumoroids generated from Patient-Derived Xenografts.* Two human colorectal tumours (TSIP#1 corresponding to LRB-0009C and TSIP#2 corresponding to IGR-0014P) from the CReMEC tumour collection *(34)* were maintained in immunocompromised mice. Animal experiments were compliant with French legislation and EU Directive 2010/63. The project received a favourable evaluation from Animal Care and Use Committee n°26 and granted French government authorisation under number 517-2015042114005883 and 8867-2017020914112908. Mice were obtained from Charles River and Gustave Roussy facility, housed and bred at the Gustave Roussy animal core facility (accreditation D94-076-11) and euthanised following endpoints validated by the Ethical Committee and the French government (Ministère de l'Enseignement Supérieur, de la Recherche et de l'Innovation). Tumoroids (TSIPs) formation from PDXs was adapted from the protocol described in *(13)*. Tumours between 1000-1500 $mm^3$ are minced and incubated in 5 to 10ml of DMEM medium containing GlutaMAX (31966-021, Gibco) supplemented with 2 mg/ml collagenase-VIII (Sigma, C2139) for 1h15 at 37°C under agitation. Tumour fragments are resuspended in 50ml of DMEM and filtered through 100µm cell strainers (EASYstrainer, 542000). Filtered tumour cells and clusters are pelleted at 800g for 10min. Pellets are further washed 4 times by adding 10ml of DMEM medium and pulse-centrifugated at 800g and 300g to collect clusters only. Clusters are cultured in suspension in DMEM medium supplemented with 10% foetal bovine serum (FBS, 10270-106, Gibco) (called "full DMEM"). They form tumoroid in 6-well ultralow attachment plates (Corning, CLS3471-24EA) after 3 days of culture. All media are supplemented with 1% penicillin and streptomycin (P/S, Gibco, 15140-122).

*Cell lines.* HT29 (ATCC® HTB-38) are grown in full DMEM. HT29-MTX (HT29-MTX-E12, 12040401, ECACC) culture medium requires the addition of 1% non-essential amino acids (NEAA, Gibco, 11140050). Cell lines are dissociated with 0.05% trypsin-EDTA. All media are supplemented with 1% penicillin and streptomycin (P/S, Gibco, 15140-122). To produce clusters from HT29 and HT29-MTX cell lines, 1.5 million cells were plated in a Petri dish, with 10ml of culture medium. Cluster formation requires 3-5 days.

*Circulating Tumour Cell lines* (CTC31, CTC44, CTC45) are a gift from Julie Pannequin and cultivated as previously described *(14)*. In brief, they are maintained in suspension as clusters in advanced DMEM-F12 (Gibco, 12634-010) supplemented with 1% GlutaMax (Invitrogen, 35050-061), 1% N2 supplement (Invitrogen, 17502-048), 20ng/ml of human EGF and 10ng/ml of human FGF-2. They are split once a week, by pelleting at 300g for 5min and incubated with Accumax for 45min at 37°C. 5ml of PBS containing 2% FBS is then added to inactivate Accumax. Clusters are filtered through a 40µm strainer, pelleted and resuspended in M12 medium.

All cell lines and clusters are cultured in a humidified incubator at 37°C under a 5% $CO_2$ atmosphere.

**Microchannels, drug incubation and cluster loading.**

*Dimensions of the channels.* Three types of microchannels were designed, with different width (w), height (h), length (l). The microchannels that were used for most migration studies have the following dimensions: h=30µm, w=60µm, l=7mm (confinement in two dimensions, one degree of freedom along X axis). The microchambers are h=30µm, w=500µm, l=7mm (providing confinement in one dimension, two degrees of freedom for migration along X and Y axes). The loading chamber's height (h=180µm) allows for quantification of clusters' displacement without confinement.

*Microchannels preparation and drug incubation.* Chips are made of a polydimethylsiloxane mixture (PDMS, Neyco, Sylgard-184 Dow Corning) 10:1 w:w with crosslinker, polymerised for at least 48h. Loading holes are made with a 1mm hole puncher. Chips are sterilised with 70% ethanol for a few minutes, dried and activated for 1min in a plasma chamber (Diener, Zepto V2, 30W) together with a glass substrate (12-well-glass-bottom plate (CellVis, P12-1.5H-N), 6-well-glass-bottom plate (MatTek, P06G-1.5-20-F) or 25-mm glass coverslip for optogenetic experiments). Channels are stuck to the activated glass before being coated for at least 30 min with an anti-adhesive reagent (0.1mg/ml pLL-g-PEG (pLL(20)-G[3.5]-PEG(2) from SuSoS) or pLL-g-PEG + 1% Pluronic F-127) or 20µg/ml rat-tail collagen-I (Corning, 354236). Channels are washed once in full DMEM and then submerged with medium for 1h to overnight. When adding drugs (Y27632, 25µM (Sigma, Y0503-5MG); Blebbistatin, 50µM (Calbiochem, 203391)), or DMSO for the control, to the medium, chips are incubated with the medium and drugs at least 3h prior to cluster loading.

*Cluster loading.* Clusters are filtered on a 70µm strainer (EASYStrainer, 542070), pelleted by a 400g pulse centrifugation and resuspended at 250 clusters/µl in full DMEM. Clusters are loaded using a 25 or 50µl syringe (Hamilton, 702SNR 22/51mm/pst3).

**Plasmids, virus production and infection**

*Plasmids*. The ARHGEF11 domain was amplified and cloned into CRY2PHR-mCherry. pCIBN(deltaNLS)-pmGFP (Addgene plasmid # 26867; http://n2t.net/addgene:26867; RRID:Addgene_26867) *(35)* and pCRY2PHR-mCherryN1 (Addgene plasmid # 26866; http://n2t.net/addgene:26866; RRID:Addgene_26866) *(35)* were a gift from Chandra Tucker. H2B-RFP and LifeAct-mCherry were gifts from the Hall lab. pRRL-Vinculin-Venus was a gift from B. Hoffman *(36)* (Addgene plasmid #111833; http://n2t.net/addgene:111833; RRID:Addgene_111833), pLV-Ftractin-mRuby3-p2A-mTurquoise-MLC-IRES-Blast was a gift from T. Meyer *(37)* (Addgene plasmid # 85146; http://n2t.net/addgene:85146; RRID:Addgene_85146) and pLentiblast-Paxillin-mTurquoise was a gift from J. Debnath *(38)* (Addgene plasmid #74206; http://n2t.net/addgene:74206; RRID:Addgene_74206). A GIPZ lentiviral shRNA transduction starter kit containing control shRNA and Talin constructs was purchased from Horizon Discovery (clone IDs (catalog number): V2LHS_56643 (RHS4430-200185645); V3LHS_366591 (RHS4430-200290073); V3LHS_366592 (RHS4430-200294817)).

*Virus production and infection.* Ectopic expression of fluorescent probes and shRNA was achieved using lentiviruses. Lentiviruses are obtained by co-transfection with the packaging vectors pMD2G (Addgene plasmid #12259; http://n2t.net/addgene:12259; RRID:Addgene 12259) and pCMVdR8,74 (Addgene plasmid #8455; http://n2t.net/addgene:8455; RRID:Addgene_8455) into HEK293T cells with the transfection reagent JetPrime (Polyplus, 114-15). Lentiviruses-containing

supernatants were collected on days 2 and 3 following transfection, concentrated by ultracentrifugation (24000g, 2h) and stored at −80 °C.
Infection was performed as described previously *(13)*. Briefly, HT29-MTX ($1x10^6$ cells) are exposed to lentiviruses in 500µl full DMEM containing 16µg/ml protamin overnight before being sorted by FACS to establish stable cell lines. Cell lines were then chosen for experiments as specified in the legends. Moreover, Vinculin-Venus-expressing HT29-MTX were used in Fig. 2, (B) and (C) (2 experiments out of 3) ; F-tractin-mRuby3/mTurquoise-MLC/Vinculin-Venus-expressing HT29-MTX were used in Fig. 3, (B) and (C) ; and for Fig. 3, (F) to (I), and fig. S5, optoRhoA HT29-MTX expressed ARHGEF11-CRY2PHR-mCherry and CIBN-GFP, and controls express CRY2PHR-mCherryN1 and CIBN-eGFP-CaaX.

**Live Imaging, microscope acquisition and optogenetic experiments.**
*Time-lapse imaging.* Timelapse bright-field imaging was done using an Olympus inverted X83 microscope with a Hamamatsu camera or a Spinning Disk CSU-W1 (Yokogawa) with a Prime 95B sCMOC camera. The latter one was also used for live fluorescence imaging.

*Optogenetics.* Clusters are incubated in the chips for at least 1 hour before imaging. Experiments were performed at 37°C in 5% $CO_2$ in a heating chamber (Pecon, Meyer Instruments, Houston, TX) placed on an inverted microscope model No. IX71 equipped with a 60× objective with NA 1.45 (Olympus, Melville, NY) and a camera ORCA-Flash4 (Hammamatsu, Japan). The microscope was controlled with the software Metamorph (Molecular Devices, Eugene, OR). Differential interference contrast (DIC) imaging was performed with a far-red filter in the illumination path to avoid CRY2 activation. Optogenetic stimulations were performed every 2-2,5 or 5min with a DMD in epi-mode (DLP Light Crafter, Texas Instruments) illuminated with a SPECTRA Light Engine (Lumencor, Beaverton, OR USA) at 440 ± 10 nm. Total Internal Reflection Fluorescence (TIRF) images were acquired using an azimuthal TIRF module (iLas2; Roper Scientific, Tucson, AZ). An automated tracking algorithm was designed in MATLAB coupled to a feedback-loop routine for the optogenetic activation.

**Immunofluorescence, antibodies, histology and immunohistochemistry**
*Immunofluorescence of TSIP#1 in collagen.* Immunofluorescence was performed on samples fixed after 3 days of incubation in collagen-1, as described previously *(13)*. Images were acquired with a SpinningDisk CSU-W1 (Yokogawa) with a Zyla sCMOC camera driven by an Olympus X83.

*Antibodies and dyes*. Primary antibodies: P5D2 (anti-integrin ß1, 1:500) was purchased from DSHB (deposited by Wayner, E.A. (DSHB Hybridoma Product P5D2)). Secondary antibodies: anti-Mouse-FITC (1:250, Jackson Immuno Research, 711-545-152). Dyes: Alexa Fluor Phalloidin 488 (1:1000, Life technology, A12379), DAPI, Alexa647-collagen-1 (coupling of the collagen was done in-house, using Invitrogen labelling kit #A20006).

*Histology and immunohistochemistry*. CRC (micropapillary histotype) obtained after chirurgical resection was formalin-fixed and paraffin embedded (FFPE) according to routine protocols. 3mm sections of FFPE samples were deparaffinised, unmasked (Ph8) and rehydrated prior Hematoxylin Eosin Saffron (HES) or immunohistochemistry.

*Immunohistochemistry.* Sections were immunostained with Ezrin (1:100, BD Biosciences, 610603) or CK20 specific mouse monoclonal antibody, (clone Ks20.8, Dako, Glostrup, Denmark). Stainings were performed with Ventana BenchMark XT immunostainer (ventana Medical Systems, Tucson, AZ) utilising UltraView DABv3 kit (Ventana). The chromogene was 3,3'-diaminobenzidine (DAB) in all the stainings.

**Image analysis**

*Analysis of clusters' displacements from bright-field time-lapse sequences*. For bright-field movies, displacement of clusters' centroid was tracked every hour using the Manual Tracking plugin in ImageJ. Clusters that were too small to be confined or dissociating during the experiment were ignored. Speed (µm/day) corresponds to the accumulated distance over 1 day (20 to 24h, and 17h for Fig.1 G and H). For observations during ≤20h, speed was extrapolated to 24h.

*Cell segmentation for cell tracking in optogenetic experiments.* Movies were analysed using custom-built routines in Fiji *(39)* and MATLAB (The MathWorks, Natick, MA). ARHGEF11-CRY2-mCherry signal was used to segment clusters by applying a gaussian filter and a threshold. Trajectories were then analysed in MATLAB, taking the displacement of the center of each segmented cluster along the microchannel.

*Persistence* is calculated as the ratio of the total displacement of the cluster monitored every 1h for 1 day, over the Euclidean distance. *Maximum instantaneous speed* is calculated as the maximum speed reached in one hour by a cluster. *Maximum migration period duration* is the longest period of continuous migration over 1px (1.07µm to 1.369µm) per hour. *Pause duration* is the length of time where a cluster stays immobile (i.e. migrates less than 1px (1.07µm to 1.369µm) per hour) between the migratory phases. Hence, one cluster can display several pauses over the imaging period. *Aspect ratio* is calculated as the clusters' length over the width of the channel (60µm).

*Analysis of Paxillin foci.* Images are treated with the Subtract background plugin, using a 2-pixels rolling ball. Threshold is then set to highlight and best separate all the foci. Characteristics of each structures is then collected using the Analyse Particles ImageJ plugin, considering particles of 10-infinite sizes.

*Analysis of nuclei movements*. Nuclei from the middle plane were imaged every 15min, manually tracked every hour and displayed using TrackMate *(40)* ImageJ plug-in. A custom-made R code was designed to subtract cluster centroid's displacement to nuclei's movement, classify the nuclei by location within the cluster (center, side 1, side 2), and represent the tracks in the cluster reference frame (Fig. S3A). As a rotation of clusters could happen in one or the other direction and could be hidden by averaging the lateral movements on several clusters, we defined Side 1 for each cluster as the cluster's side going most rearward (or less forward) in average, while the other side was called side 2.

*Analysis of MLC and actin polarisation*. Direction of migration for each cluster is determined by tracking its movement for at least 2h prior to fluorescence imaging. Myosin and actin localisation are determined in the middle section of the sphere using a spinning disk fluorescence microscope. 20-pixel Subtract background is performed before using a 15 pixels wide line scan of the cortex to measure the intensity of myosin and actin signal in each half of the cluster. The intensity in each half is normalised to the perimeter of the line scan to calculate the rear to front ratio.

**Western blot**.
Western blots were conducted as described previously *(13)*, with PVDF membranes (GE Healthcare). Primary antibody was anti-Talin 1 antibody (ab71333, Abcam) and secondary antibody was anti-rabbit, HRP-linked (7074, Cell Signaling).

**Statistics and reproducibility**
Normality or lognormality of all data distribution was tested using the Kolmogorov-Smirnov, d'Agostino-Pearson or Shapiro-Wilk test in Prism 8 (GraphPad). When distributions are best fitted by a lognormal distribution, statistical tests were performed on log-transformed data. Significance for datasets displaying normal distributions were calculated in Prism with unpaired two-tailed Student's t test or one-way analysis of variance (ANOVA) and Holm-Sidak's multiple comparison test when comparing more than two conditions. Significance for non-normal distributed datasets were calculated in Prism using a Mann-Whitney test or a Kruskal-Wallis test and Dunn's multiple comparison test when comparing more than two conditions. Significance for non-normal distributed paired datasets (Fig. 2K) were calculated in Prism using Friedman test and Dunn's multiple comparison test when comparing more than two conditions. P values of statistical significance are represented as ****P <0.0001, ***P <0.001, **P <0.01, *P <0.05, ns, not significant. The exact value is indicated when possible. n numbers are indicated in the figure legends as well as the N number of independent experiments. Violin plots display the whole population of clusters, median (dashed grey line) and quartiles (dotted grey lines) *(41)*. The coloured dots represent the mean of each independent experiments and the black line is the mean of all experiments for each condition. Experiments were performed independently for each cell lines and for each individual graph, coloured dots for the same cell line refer to the same experiment.

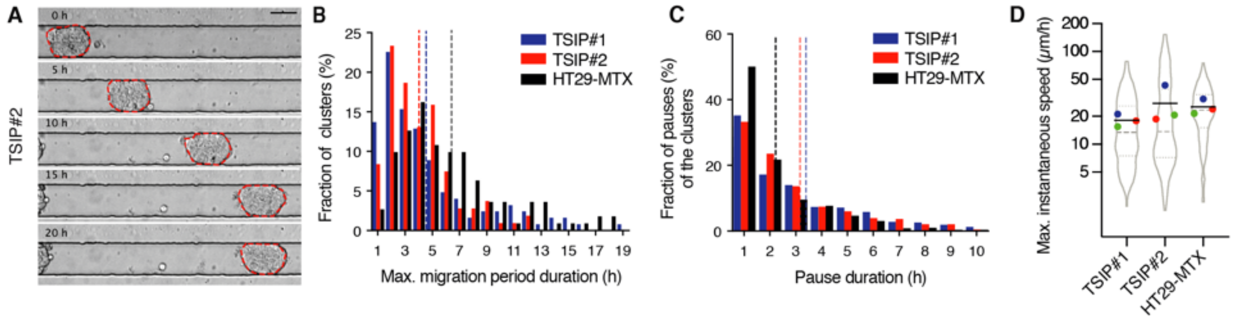


**Fig. S1.**

**Characterisation of clusters' migration.** (**A**) Time-lapse sequences of TSIP#2 migration every 5h migrating in a PEG-coated microchannel. Scale bar, 50$\mu$m. (**B**) Histogram of the longer period of consecutive migration achieved by each cluster (maximum (Max.) migration period duration) migrating one day in PEG-coated microchannels. (**C**) Histogram of the duration of pauses occurring during clusters' migration during one day in PEG-coated microchannels. For (B) and (C), dashed lines: mean. (**D**) Maximum instantaneous speed of clusters migrating one day in PEG-coated microchannels, log2-scale. For (B) to (D), n=107 to 124 clusters from 3 independent experiments. Violin plot is described in Fig. 1 legend.

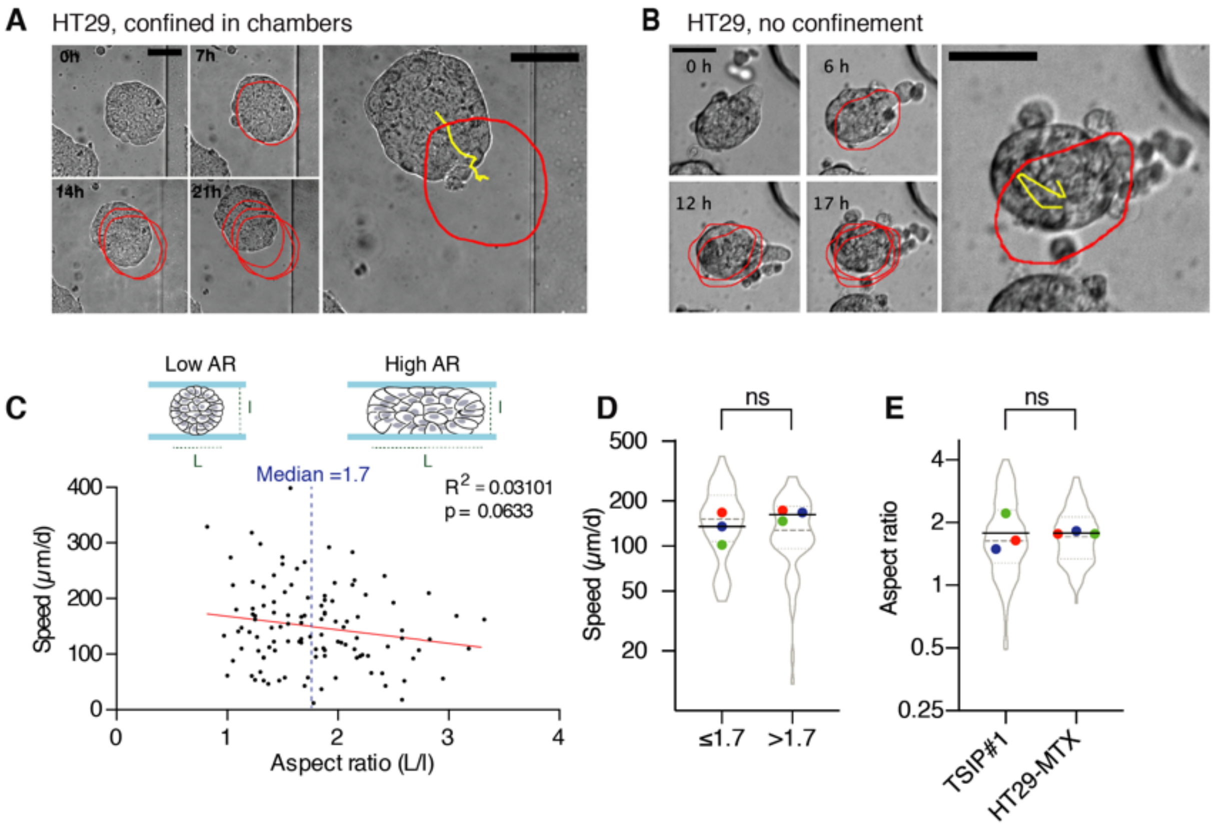


**Fig. S2.**

**Confinement increases the persistence of cluster migration in non-adhesive microchannels, independently of their size.** (**A** and **B**) Representative time-lapse sequences of HT29 displacement, confined in microchambers (A) and in loading chambers (B, no confinement), coated with PEG+F127. Red circles: consecutive positions (every 6 or 7h); yellow: track of the cluster's centroid monitored every hour for one day. Scale bars, 50$\mu$m. (**C**) Speed and aspect ratio (AR) of HT29-MTX clusters migrating one day in PEG-coated microchannels. n=112 clusters from 3 independent experiments. Line: linear regression. (**D**) Comparison of the speed of HT29-MTX clusters migrating one day in PEG-coated microchannels, depending on their aspect ratio (relative to median AR=1.7, illustrated in (C)), log2-scale. n=112 clusters from 3 independent experiments (Mann-Whitney test). (**E**) Aspect ratio of clusters migrating one day in PEG-coated microchannels, log2-scale. n=112 or 162 clusters from 3 independent experiments (two-tailed Student's t test). ns, not significant. Violin plots are described in Fig. 1 legend and in (D), coloured dots refer to the same experiment. All data represented as violin plots are from N=3 independent experiments.

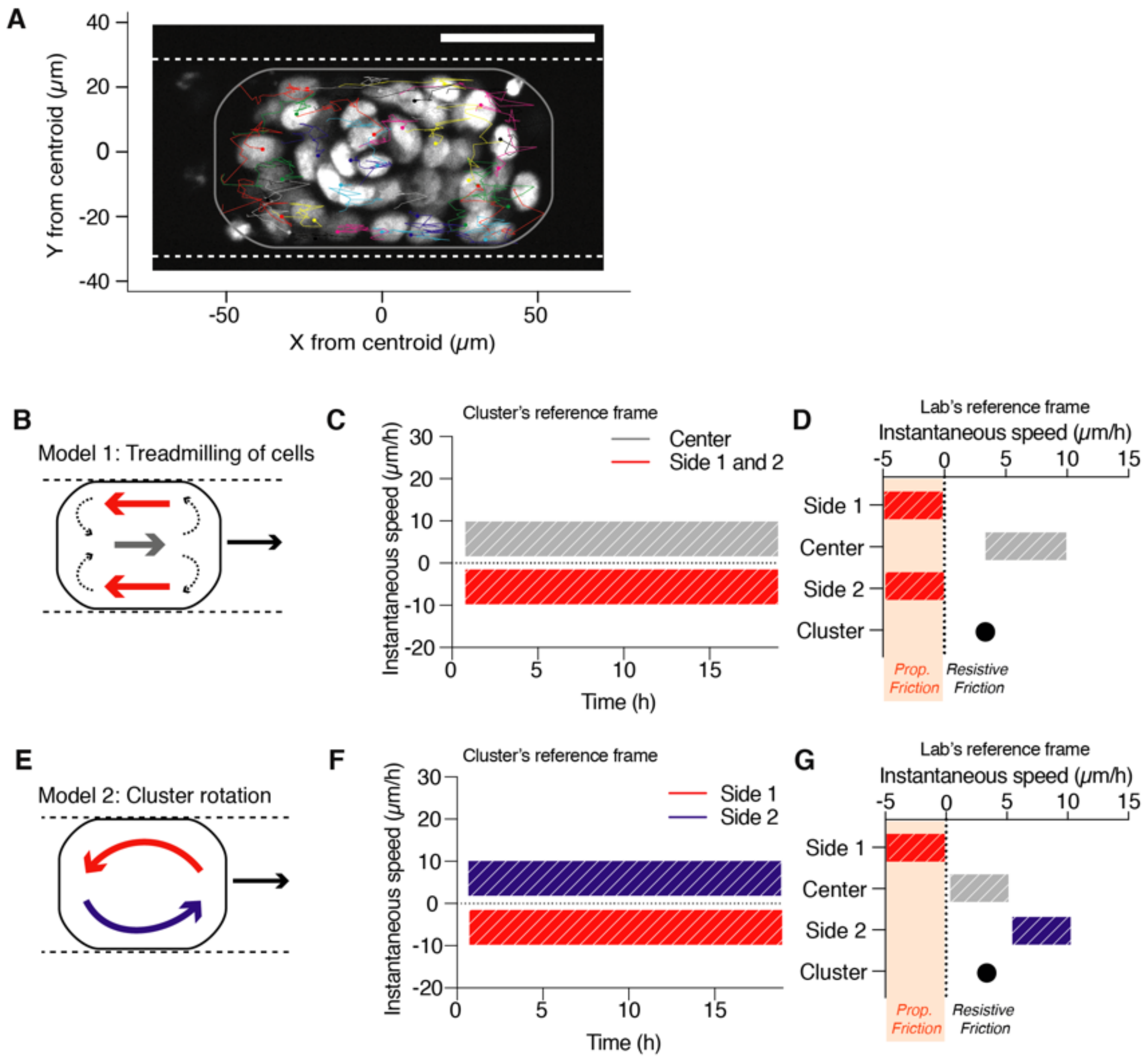


**Fig. S3.**

**Adhesion-independent collective migration is not mediated by a cell flow.** (**A**) Representative tracks of nuclei in the cluster's reference frame, migrating in a PEG-coated microchannel over one day (same cluster as in Fig. 2H). (**B** to **G**) Hypothetical models, corresponding to Fig. 2, (I) to (K). Schematic representation of cell treadmilling (Model 1, B) or rotation of the whole cluster (Model 2, E) generating propulsive friction with microchannel walls. (C and F) Expected range (hatched rectangles) of instantaneous speed of the cells present in each zone, in the cluster's reference frame. (D and G) Expected range (hatched rectangles) of mean instantaneous speed of cells in contact with the microchannel walls, considering the same mean instantaneous speed of the cluster as in Fig. 2K, lab's reference frame. Orange zone corresponds to cell speeds ≤0 capable of generating a propulsive (Prop.) friction against the microchannel walls

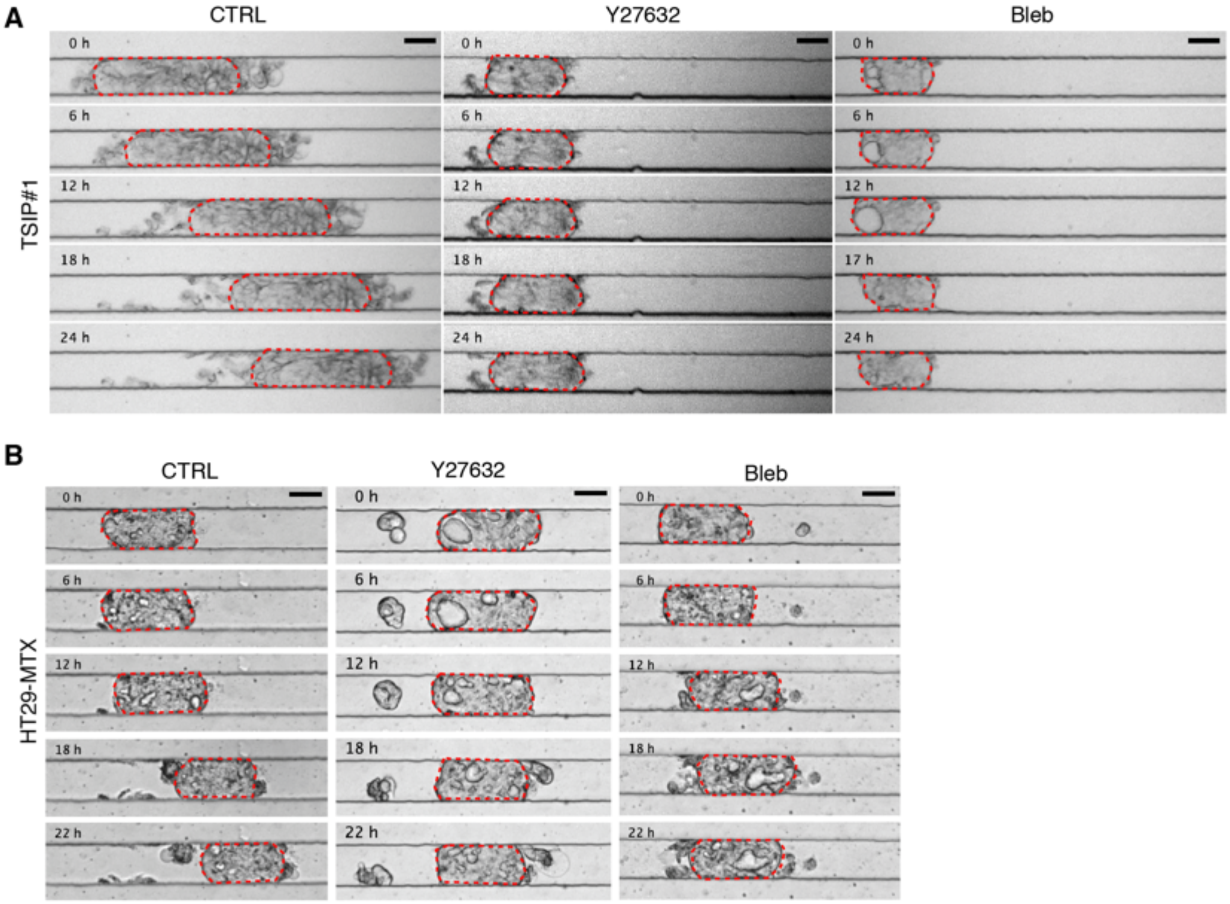


**Fig. S4.**

**Inhibition of acto-myosin contractility prevents focal adhesion-independent collective migration.** Time-lapse sequences of TSIP#1 (**A**) and HT29-MTX (**B**) displacement in PEG-coated microchannels over a day. (Left) control condition, (middle) after Y27632 (25$\mu$M) treatment, (right) after Blebbistatin (50mM) treatment. Scale bar, 50$\mu$m.

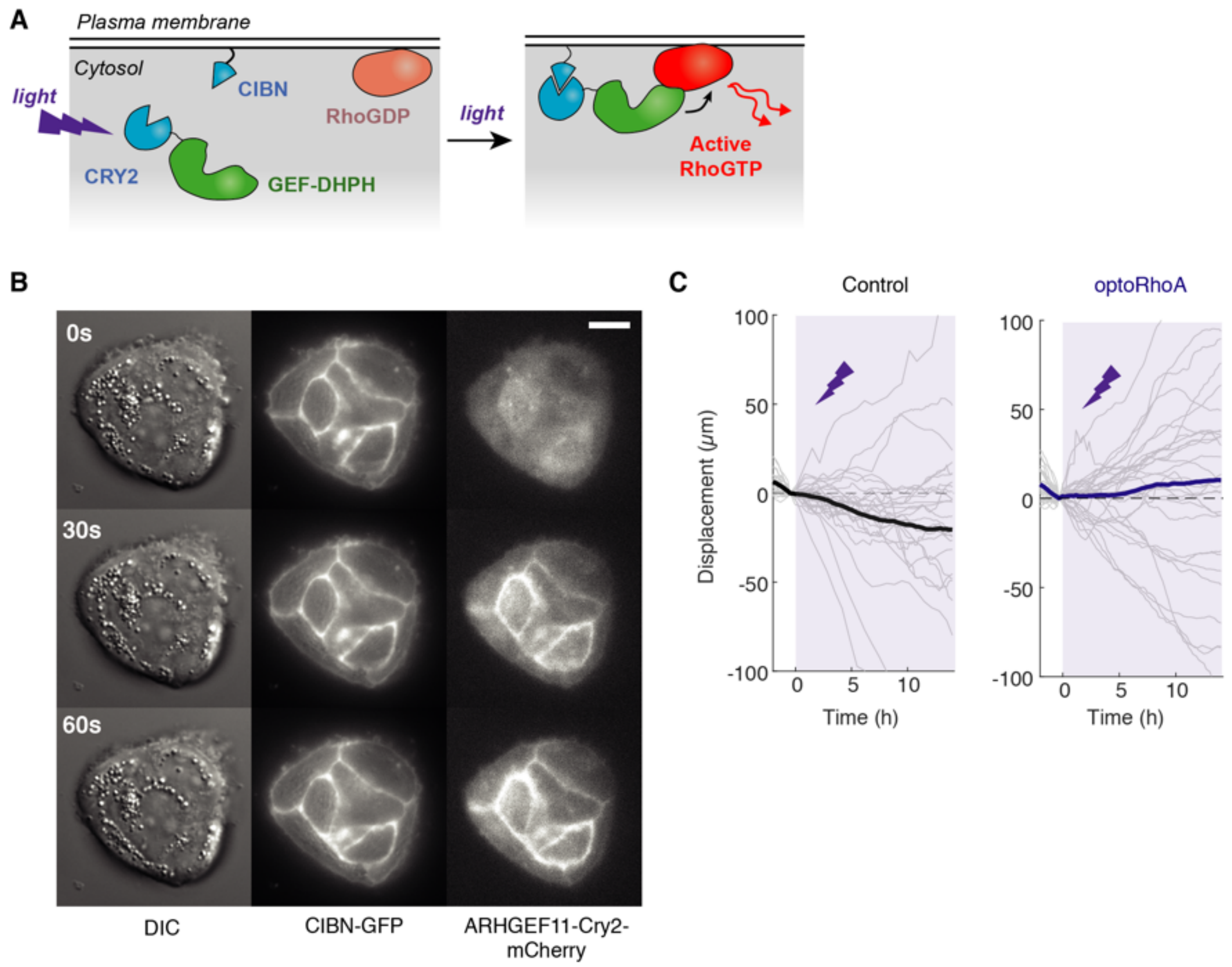


**Fig. S5.**

**Activation of RhoA induces cluster migration and dictates directionality.** (**A**) Schematic of the molecular effect of light activation in optoRhoA cells. (**B**) Global activation of optoRhoA cell lines. ARHGEF11-Cry2-mCherry and CIBN-GFP are observed in Total Internal Reflection Fluorescence (TIRF). Activation every 30s with 488nm-laser. Scale bar, 10$\mu$m. (**C**) Displacement of clusters (grey) before (-2h<t<0h) and after (0h<t<15h) optogenetic activation of optoRhoA stably expressing cells and control cells. Means are represented in bold. Purple zone: optogenetic activation. n=27 for optoRhoA from 3 independent experiments.

**Movie S1.**

TSIP#1 (top) and TSIP#2 (bottom) clusters migrate collectively in PEG-coated microchannels. Movie was recorded every hour over 24 hours. Scale bars, 50μm.

**Movie S2.**

HT29-MTX clusters migrate in PEG-coated microchannels. Movie was recorded every hour over 20 hours. Scale bar, 50μm.

**Movie S3.**

HT29-MTX clusters stably expressing Paxillin-mTurquoise in a collagen-I-coated microchannel (left) or in a PEG-coated microchannel (right). Movie was recorded every 3 min over 2.5 hours. Scale bar, 30μm.

**Movie S4.**

HT29 cluster stably expressing mCherry-H2B migrating in a PEG-coated microchannel. Movie was recorded every 15min over 24 hours. Scale bar, 50μm.

**Movie S5.**

HT29-MTX clusters stably expressing mTurquoise-MLC static (left) or migrating (right) in PEG+F127-coated microchannels. Movie was recorded every 5 min over 3 hours. Scale bar, 30μm.

**Movie S6.**

OptoRhoA HT29-MTX cluster migrating in a PEG-coated microchannel before and after optogenetic activation. Optogenetic activation, represented in blue, starts at 0min. Movie was recorded every 10min over 14h hours. Dark grey lines represent microchannel walls. Scale bar, 20$\mu$m.

**Movie S7.**

CRY2PHR-mCherryN1/CIBN-eGFP-CaaX-expressing HT29-MTX cluster (control for optoRhoA clusters) migrating in a PEG-coated microchannel before and after optogenetic activation. Optogenetic activation, represented in blue, starts at 0min. Movie was recorded every 10min over 15h hours. Dark grey lines represent microchannel walls. Scale bar, 20μm.
